# The Improbable Journeys of Epiphytic Plants Across The Andes: Historical Biogeography of *Cycnoches* (Catasetinae, Orchidaceae)

**DOI:** 10.1101/106393

**Authors:** Oscar Alejandro Pérez-Escobar, Marc Gottschling, Guillaume Chomicki, Fabien L. Condamine, Bente Klitgård, Emerson Pansarin, Günter Gerlach

## Abstract

The Andean uplift is one of the major orographic events in the New World and has impacted considerably the diversification of numerous Neotropical organisms. Despite its importance for biogeography, the specific role of mountain ranges as a dispersal barrier between South and Central American lowland plant lineages is still poorly understood. The swan orchids (*Cycnoches*) comprise *ca* 34 epiphytic species distributed in lowland and pre-montane forests of Central and South America. Here, we study the historical biogeography of *Cycnoches* to better understand the impact of the Andean uplift on the diversification of Neotropical lowland plant lineages. Using novel molecular sequences (five nuclear and plastid regions) and twelve biogeographic models with and without founder-event speciation, we infer that the most recent common ancestor of *Cycnoches* may have originated in Amazonia *ca* 5 Mya. The first colonization of Central America occurred from a direct migration event from Amazonia, and multiple bidirectional trans-Andean migrations between Amazonia and Central America took place subsequently. Notably, such biological exchange occurred well after major mountain building periods. The Andes have not acted as an impassable barrier for epiphytic lowland lineages such as orchids having a great potential for effortless dispersal because of the very light, anemochorous seeds.

## Introduction

Neotropical landscape and biodiversity have long drawn the attention of naturalists ^1,2^. The tropical Andes are of particular interest as the world’s premier biodiversity hotspot, with both an extraordinary species richness and a remarkable level of endemism^3– 5^. The combination of molecular phylogenies with species distributions and the fossil record has uncovered different biotic and abiotic factors that fostered diversification in the Neotropics^6–  10^. However, biogeographical studies applying modern phylogenetic methods are only available for few Neotropical plant clades (e.g., *Begonia*^11^, *Cyathostegia*^12^*, Heliotropium*^13^, *Lupinus*^14,15^, palms^16^, Rubiaceae^6^). These studies have generally demonstrated the importance of geological processes such as mountain building and establishing the Isthmus of Panama for the diversification of Neotropical plants^17^.

One of the most biologically relevant abiotic processes in the diverse geological history of the Americas is the rise of the Andes ^7,13,18^. Andean mountain building was driven by plate tectonic re-adjustments that started during the Palaeogene and continued until the Pliocene ^7^. The fossil record (e.g., palynological^3,19^ and geological data: isotope measurements^20^, sediment loads, apatite fission-track data^7^) collectively indicate that the Andean uplift was a partially constant process punctuated by periods of intensified mountain building. Two of the most intense uplift periods occurred around 12 (mid-Miocene) and 4.5 million years ago (Mya; early Pliocene^7^). During these periods, the Northern Andes reached elevations as high as 4500 m in the Pliocene, whereas the Central Andes already peaked an altitude of 4500 m during the mid-Miocene ^7,21^.

Newly formed mountain ranges may had a strong impact on the adjacent Amazonian landscapes and the inhabiting organisms due to the transformation of its drainage systems^22^. The Andes also influenced local and regional climates by forming the only barrier to atmospheric circulation in the Neotropics^23,24^. The rise of the Andes led to the formation of island-like habitats and of local microclimates and soil conditions that eventually fostered species diversification^14,25,26^. At the same time, the Andes provided physical and/or ecological barriers to species dispersal and migration. For instance, the efficiency of the Northern Andes as migration barrier is shown for Andean centred woody species of *Crematosperma*, *Klarobelia*, *Malmea*, and *Mosannona* (Annonaceae^27^). Propagules are dispersed by animals in these lineages^28^, and none of their constituent species occur in both east and west of the Andes mountain range^27^.

Recent phylogenetic studies provide solid evidence for the important role of Andean uplift in diversification of highland-dwelling plant groups (e.g., *Lupinus*^14^; *Bartsia*^29^; centropogonid Campanulaceae^9^). However, the impact of such orographic processes for the lowland flora is still poorly understood^4,6,30^. Thus, the question remains whether Andean uplift has indeed been an abiotic barrier to migration for epiphytic lineages such as lowland orchids and bromeliads, both being important components of Neotropical forests.

Epiphytic diversity is dramatically greater in the Neotropics than in any other tropical region around the world ^31,32^, being twice as high than, for instance, in Australasia ^33–  36^. Several traits shared by Neotropical epiphytic taxa, and related to their reproductive biology, may explain such overwhelming differences in diversity. One of the most prominent shared traits is the lightness and the very small size of the propagules, occasionally with highly elaborated surfaces (e.g., bromeliads, ferns, orchids, *Pinguicula*^33^). The potential of dust-like seeds for longer distance dispersals by wind might be much greater compared to other plants with propagules locally dispersed by animals (e.g., Araceae^37^). Nonetheless, it remains largely unknown whether lowland epiphyte lineages have been able to disperse across geological barriers such as the Andes. Well-sampled phylogenies for epiphyte clades have been lacking to address this issue.

Several wind-dispersed plant lineages (e.g., *Begonia*^11^; bromeliads^38^) span across the Neotropical region, many of which are restricted to lowland elevations. One example is the orchid tribe Cymbidieae comprising *ca* 3900 species that are mostly distributed in the Neotropics (but with few representatives in the Old World Tropics^39^). To Neotropical Cymbidieae belong the swan orchids (*Cycnoches*) that are known for the striking sexual dimorphisms^40^ and pollination syndrome^41^. Molecular phylogenetics and morphological studies conducted to date confirm the inclusion of *Cycnoches* within Catasetinae^42–44^ and its sister group relationship to *Mormodes*^40,45,46^.

*Cycnoches* encompasses 34 species of epiphytes^47^ that are distributed from Southern Mexico to Central Brazil and Bolivia. Its species are best represented in the Amazonian forests of Brazil, Ecuador, and Peru, but also in the Llanos and Caribbean regions of Colombia and Venezuela^47,48^. They commonly inhabit lowland tropical wet forests, ranging from 0 to 800 m. *Cycnoches* species are pollinated by male euglossine bees, which collect volatile compounds produced by flowers but also from other sources (e.g., rotten wood, faeces^41^). They have rather a restricted geographical range, and most of them are distributed in single biogeographical area. Nevertheless, one species (*C.chlorochilon*) is distributed in both sides of the Northern Andes^47^.

Because of the striking sexual system they have evolved^40^, swan orchids have long attracted the attention of several prominent botanists including^49^. Despite this long interest, previous phylogenetic studies have only been included up to three species of *Cycnoches* probably because of their scarceness both in the wild and in herbarium collections^42,43,45,46,50^. Moreover, the lack of a solid and well-sampled phylogeny of *Cycnoches* has precluded addressing specific questions concerning the role of Andean uplift in their biogeographical history. The narrow geographical distribution of almost all extant *Cycnoches* species and their restricted habitat preference given (i.e., lowland wet forests, see above), we expect the swan orchids diversification to be strongly influenced by the Andean uplift. In particular, we hypothesize that Andean uplift was an impassable, isolative, barrier for *Cycnoches*, as already reported for other plant lineages. To test this hypothesis, we generated a strongly supported 5-loci phylogeny, sampling 23 out of 34 accepted *Cycnoches* species and comprising its known diversity and distribution, and performed models of biogeographical analyses.

## Results

### Phylogeny of Cycnoches

Our phylogeny comprised 23 out of 34 accepted species of *Cycnoches*. Table S5 provides detailed alignment descriptions. The concatenated ‘nuc’ alignment was 2395 bp in length and included 310 parsimony informative sites, and the concatenated ‘cp’ alignment was 2419 bp and comprised 171 parsimony informative positions. Independently derived ‘nuc’ and ‘cp’ phylogenies revealed topologies with conflicting and highly supported phylogenetic placements. PACo analysis revealed 22 potential outlier OTUs (see below; Figs S1–S2) belonging to *Cycnoches* (majority of 19), *Dressleria*, and *Mormodes*. After inspection of the potentially confliting OTUs placement in ‘nuc’ and ‘cp’ phylogenies (Fig. S2), 20 outliers were confirmed as conflicting terminals. Thus, only two terminals (i.e., *Dressleria severiniana* and one representative of *C. lehmannii*) were misclassified by PACo as conflicting (see Appemdix S1 for a detailed explanation on outlier handling).

Within *Cycnoches*, the ‘nuc’ phylogeny recovered three maximally supported clades (A, B, C; Fig. 1). Clade A included all sequenced accessions of *Cycnoches hagii* and was recovered as sister group of the remaining species of *Cycnoches* comprising clades B and C. By contrast, the ‘cp’ phylogeny showed two main maximally supported clades (namely I and II), each including a set of intermingled species from clades A, B, and C. The best ML tree inferred from non-conflicting, concatenated ‘nuc’ and ‘cp’ datasets showing the internal phylogenetic relationships of *Cycnoches* is presented in Figure 2. Virtually, all internal nodes of the backbone phylogeny were highly, if not maximally supported by MLBS and PP values. *Cycnoches* segregated into three main lineages (clades A, B and C). Clade A (i.e., all specimens of *C. haagii*) was sister group of the remaining species of *Cycnoches* clustering in clades B and C.

**Figure 1.**
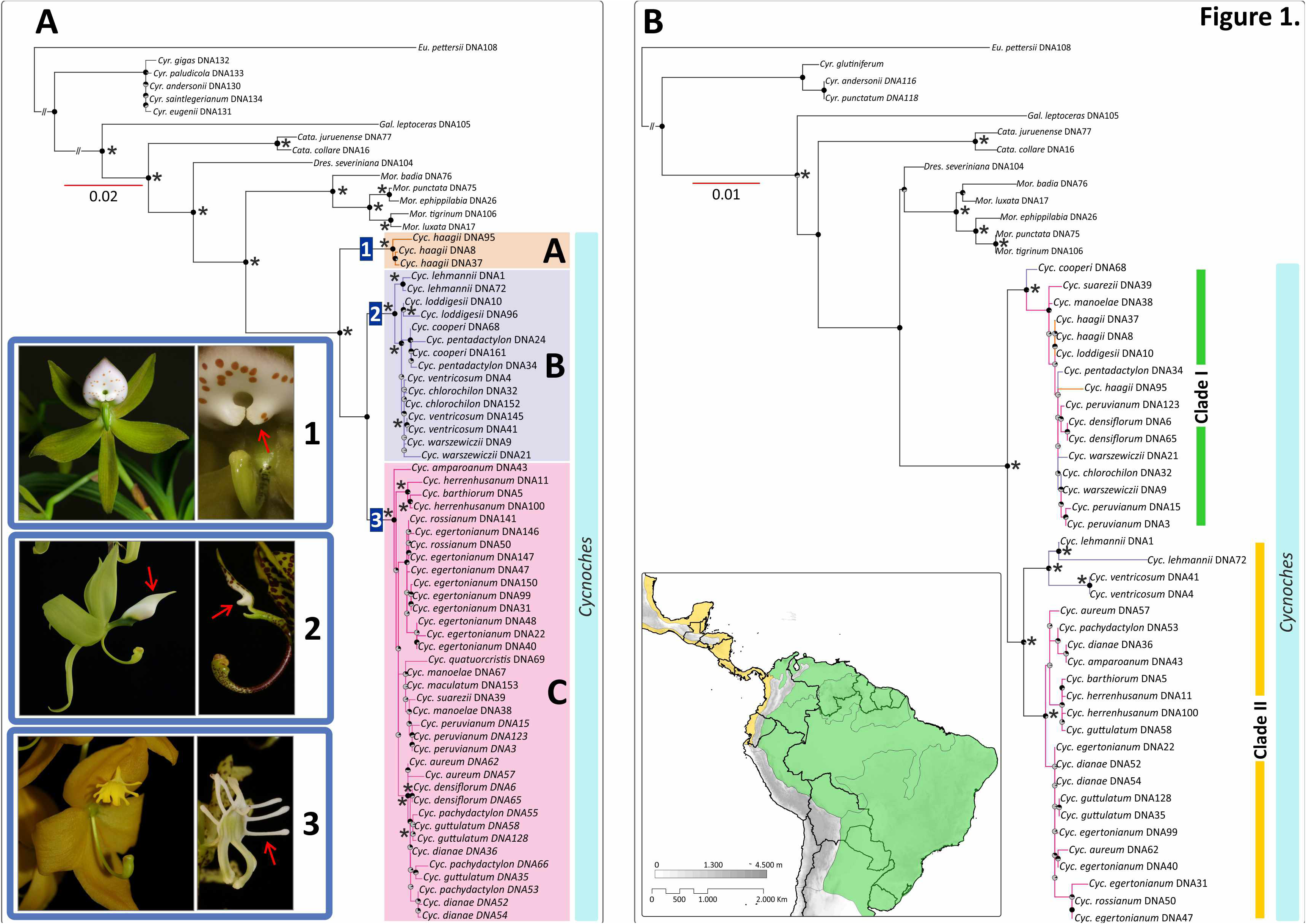
Best scoring, ML tree of *Cycnoches* obtained from concatenated **A)** nuclear ETS,ITS, *Xdh* and **B)** plastid *trn*S-G, *ycf*1 loci. Node charts indicate Bootstrap Support (BS > 75), in where fully red diagrams indicate BS of 100. Numbers at nodes indicate Bayesian Posterior Probability (PP > 0.95). Representatives of each clade are shown in pictures: **1)***C.haagii*; **2)***C. ventricosum* (left) and *C. pentadactylon* (right); **3)***C. herrenhusanum* (left)and *C. peruvianum* (right). The red arrows indicate autapomorphies of every clade: in 1) fleshy oblong, curved pair set of calli located towards the base of the labellum, 2) labellum blade entire to 4-lobed, and 3) labellum with 8-10 marginal projections. Geopolitical boundaries map generated by ArcMAP software (http://www.esri.com), using layers freely available at http://www.diva-gis.org/Data. Photos: O. Pérez & G. Gerlach.

**Figure 2.**
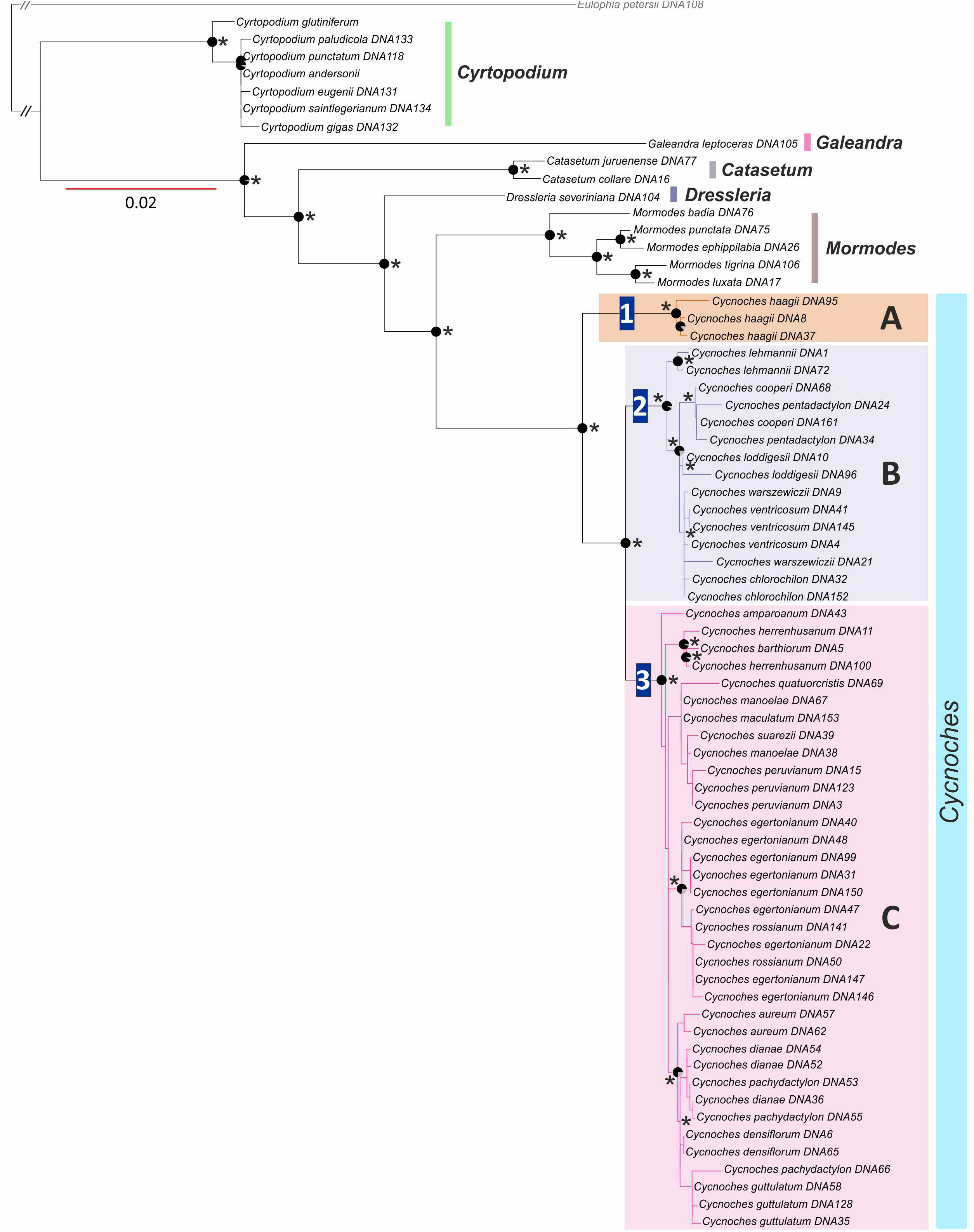
Best scoring, ML tree of *Cycnoches* obtained from concatenated, non-conflicting‘nuc’ ETS, ITS, and *Xdh* and ‘cp’ *trn*S–*trn*G and *ycf*1 loci. Node charts indicate Bootstrap Support (BS > 75), in where fully red diagrams indicate BS of 100. Numbers at nodes indicate Bayesian Posterior Probability (PP > 0.95).

### Molecular dating of Cycnoches

Comparison of marginal likelihood estimates (MLE) of tree priors and clock models revealed that an uncorrelated molecular clock combined to a birth-death tree prior with incomplete sampling best fitted our data (MLE of –14244.47; Table 1). Nevertheless, Bayes factors (BF) clearly rejected the strict clock models (BF = 72, 45, and 40 for all tree priors), but did not provide strong evidence to support an incomplete sampling birth-death model *vs* a standard birth-death model (BF=1.08; Table 1). Analysis of the log file produced by dating analyses under the relaxed clock and tree models are also shown in Table 1. Overall, they yielded CV values between 0.35 and 0.42 (indicating there was among branch rate heterogeneity, which argued for the use of a relaxed molecular clock). Therefore, results obtained from a dated phylogeny with relaxed molecular clock and birth-death standard speciation model (Table 1) are presented only (Figs 3, S5).

**Table 1.** Marginal likelihoods of tree speciation models for relaxed and strict molecular clocks.

|  | **Tree prior** | **MLE**<br>**(SS)** | **CV** | **Yule** | **BD** | **BD**<br>**sampling** | **Yule** | **BD** | **BD**<br>**sampling** |
| --- | --- | --- | --- | --- | --- | --- | --- | --- | --- |
| **Strict clock** | **Yule** | -14280,93 | - | - |  |  |  |  |  |
|  | **BD** | -14267,36 | - | 27,14 | - |  |  |  |  |
|  | **BD**<br>**sampling** | -14264,73 | - | 32,4 | 5,26 | - |  |  |  |
| **Relaxed clock** | **Yule** | -14259,53 | 0,42 | 42,8 | 15,66 | 10,4 | - |  |  |
|  | **BD** | -14245,01 | 0,35 | 71,84 | 44,7 | 39,44 | 29,04 | - |  |
|  | **BD**<br>**sampling** | -14244,47 | 0,35 | 72,92 | 45,78 | 40,52 | 30,12 | 1,08 | - |

A chronogram showing absolute ages estimated under a relaxed clock is presented in Figure 3 (see also Fig. S5 for the 95% confidence intervals) and shows that *Cycnoches* and *Mormodes* shared a common ancestor during the beginning of the late Miocene (9.1 Mya ±3). Diversification of *Cycnoches* took place around 5 Mya ±2 at end of the late Miocene. The split between *Cycnoches* clades B and C occurred during late Pliocene (3 Mya ±2), whereas both clades were estimated to be of Pleistocene ages (1.2 and 1.6 Mya ± 1, respectively).

**Figure 3.**
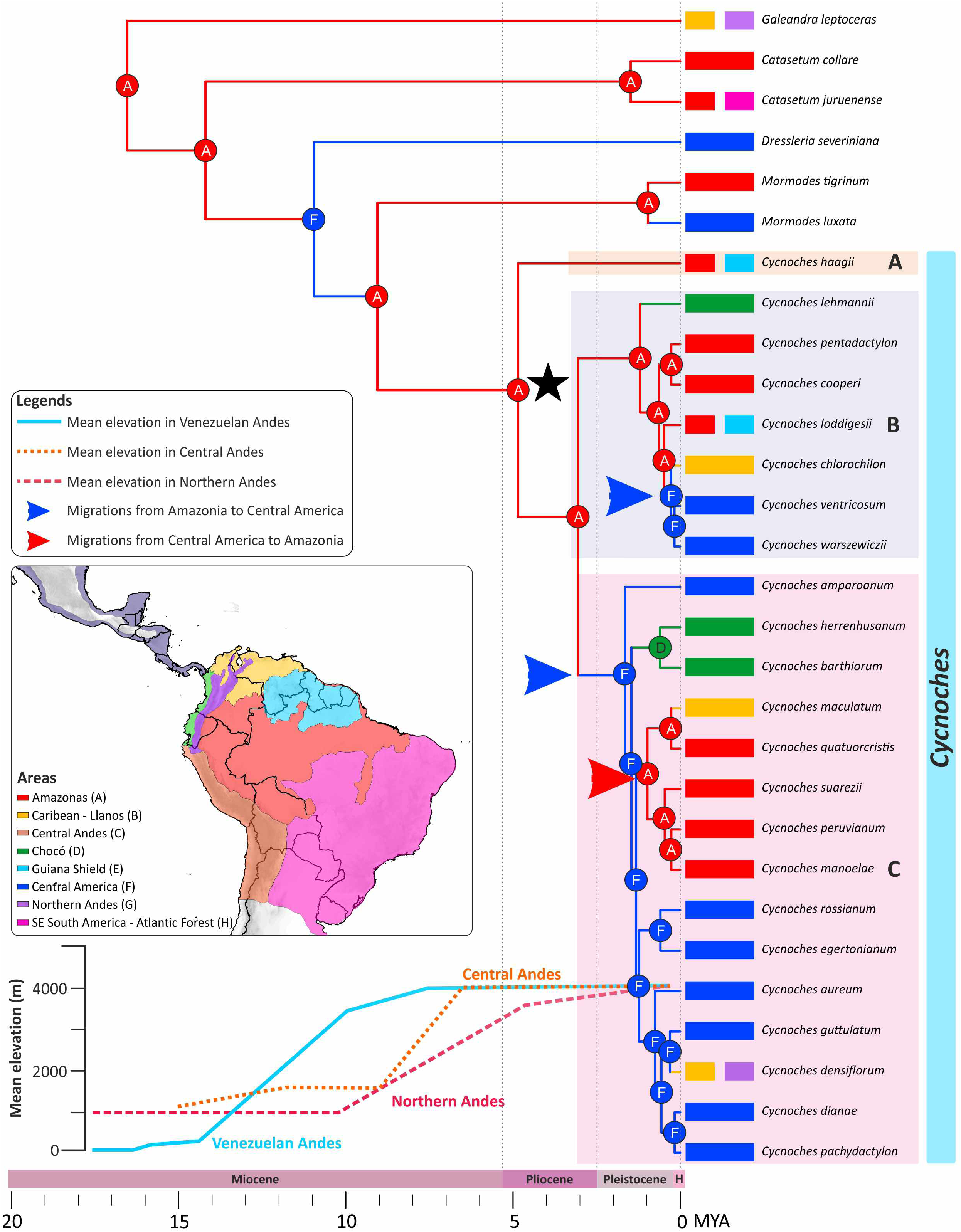
Chronogram for *Cycnoches* obtained under a relaxed clock model, applied to anon-conflicting, concatenated ‘nuc’ (ITS, ETS, *Xdh*) and ‘cp’ loci (*trn*S–*trn*G, *ycf1*). Age estimates, including maximum and minimum intervals for all nodes, are provided in Figure S5. Time scale is provided in million years ago (Mya). Node charts correspond to ancestral areas estimated under the BayArea-Like* model, including founder event process (*J*). Pink, orange and blue lines indicate mean elevations (m) on Colombian and Venezuelan Northern Andes, respectively (adapted from Hoorn et al., 2010). (Inset) Coded areas used for biogeographical analysis. Geopolitical boundaries map generated by ArcMAP software (http://www.esri.com). Political divisions and elevation data from DIVA-GIS (http://www.diva-gis.org/gdata).

### Ancestral area estimations

Table 2 provides ML statistics for the biogeographical models as inferred in BioGeoBEARS. The best fitting parametric model was the BAYAREA star, including the founder-event speciation. This model revealed a geographical origin in Amazonia as the most likely ancestral area of *Cycnoches* (Figs 3, S6). The MRCA of clades B and C was reconstructed to have inhabited in Amazonia region. The MRCA of clade C occurred in Central America, whereas the MRCA of clade B inhabited in Amazonia region.

**Table 2.** Likelihood of biogeographic models as implemented in DEC, DIVA-like andBayArea-like.

| **Model** | **LnL** | **AIC** | **AICc** | **AIC weighted** |
| --- | --- | --- | --- | --- |
| DEC | -119,3054256 | 242,6108513 | 242,9744877 | 1,06E-08 |
| DEC+J | -110,4860209 | 226,9720418 | 227,7220418 | 2,63E-05 |
| DEC* | -106,2412542 | 216,4825085 | 216,8461448 | 0,004984487 |
| DEC+J* | -103,9600773 | 213,9201546 | 214,6701546 | 0,017948515 |
| DIVA | -125,2827663 | 254,5655327 | 254,929169 | 2,68E-11 |
| DIVA+J | -123,1895028 | 252,3790055 | 253,1290055 | 7,99E-11 |
| DIVA* | -104,1616852 | 212,3233704 | 212,6870068 | 0,039880977 |
| DIVA+J* | -104,9727468 | 215,9454937 | 216,6954937 | 0,006519762 |
| BAYAREA | -131,2239925 | 266,4479849 | 266,8116213 | 7,04E-14 |
| BAYAREA+J | -113,2503658 | 232,5007317 | 233,2507317 | 1,66E-06 |
| BAYAREA* | -119,7393191 | 243,4786382 | 243,8422745 | 6,85E-09 |
| **BAYAREA+J\*** | **-100,011714** | **206,0234279** | **206,7734279** | **0,930638293** |

Three independent trans-Andean migration events between Amazonia and Central America could be identified. The first migration from Amazonia to Central America took place towards the late Pliocene (±1 Mya), after the divergence of MRCAs of clades B and C. A second migration from Central America to Amazonia took place around mid-Pleistocene (±1 Mya) by the MRCA of *C. maculatum*, *C. manoelae*, *C.peruvianum*, *C. quatuorcristis*, and *C. suarezii*. Last biotical exchange from Amazonia to Central America was dated to late Pleistocene (±0.5 Mya) with the MRCA of *C.chlorchilon*, *C. ventricosum*, and *C. warzsewiczii*.

## Discussion

### Influence of Andean orogeny on the biogeography of a Neotropical epiphyte group

Our study provides a solid phylogenetic framework, divergence times, and ancestral areas estimation for *Cycnoches*. Central America has been considered the most likely region of *Cycnoches* origin, probably because of its locally elevated species richness as compared to other areas in the Neotropics (Romero & Gerlach, *in press*). However, this evolutionary scenario is rejected by our analyses, which instead support a South American origin of *Cycnoches* in the late Miocene (ca. 5 Mya ±2; Fig. 3, Fig. S6). The ancestral area estimated at *Cycnoches* MRCA largely reflects the current distribution of early diverging lineages such as *C. haagii*, a species today distributed in Amazonia and the Guiana Shield. Thus, the early diversification of *Cycnoches* has taken place well after the most intense mountain building events of the Northern and Central Andes ca. 12 and 10 Mya, respectively^7,52 –54^.

Our results indicate a less important role of the Andes as a biogeographic barrier for the diversification of *Cycnoches*. Two migrations from Amazonia to Central America and one reverse colonisation event back imply vigorous ancient dispersals across the Andes during last 5 million years (Figs 3). During the early Pliocene (i.e., the period when *Cycnoches* has started to diversify), the Northern Andes of Colombia and Venezuela have already reached elevations up to at least 3000 m (Hoorn *et al.*, 2010). Moreover, three migrations from Amazonia to Central America and back, respectively, took place ~2 Mya, when the Northern and Central Andes already peaked at a mean elevation of 4000 m (see mean Andean elevation displayed in the inset of Fig. 3).

Similar trans-Andean migrations are reported for bromeliads and ferns. In Bromeliaceae (subfamily Bromelioideae), direct migrations from the Brazilian Shield towards Central America took place around 7 Mya^38^, a period where Northern Andes reached a palaeoelevation of ~2000 m^7^. Furthermore, biotical exchanges between South and Central America have also been reported in the fern complex *Jamesonia-Eriosorus* (Pteridaceae)^30^. Here, migrants from the Brazilian coast colonised and further established in Central Andes, and from there subsequently migrated towards Central America during the late Pleistocene^55^.

Impermeability of the Andes as a barrier for reproductive isolation does not appear as important for Neotropical epiphyte lineages such as *Cycnoches*. Our results also point to a rapid colonization of the mainland Neotropics (i.e., colonization of Central America, Amazonia, and Choco in ~5 million yr.). Such biotic invasion may be related to the biology of the group having anemochorous seed dispersal and an epiphytic habit. Most orchid seeds are characterized by their minute size and reduced weight, as well as by their elaborated seed coats^56,57^. These traits allow them to easily remain airborne for extended periods of time and to travel over long distances^57^. *Cycnoches* species might have good dispersal potential as well because of their seeds, which are between 100–300 µm long, 50–60 µm wide, and about 3.6 µg weight^56,58^.

Empirical evidence strongly supports short mean dispersal distances (e.g., 4–5 m) of seeds from terrestrial orchids such as *Orchis purpurea*^59^, yet there are no experimental studies supporting putatively longer dispersal of epiphytic orchid seeds. Nevertheless, disjunctive distributions of several orchid taxa occurring, for instance, in remote islands (e.g., *Anacamptis pyramidalis* growing in Aldabra island and Madagascar^60^) require dispersals over longer ranges as explanation. Authors have already reported disjunctive distributions of several orchid species resulting from putative long-distance dispersals^61,62^. However, to the best of our knowledge, no study has yet reported ancient orchid long-distance dispersal across a geographical barrier such as the Andes.

### Phylogenetic conflict between nuclear and plastid phylogenies in Cycnoches

Our study brings important insights for the species relationships within *Cycnoches*. Previous phylogenetic studies about Catasetinae have included not more than three species of *Cycnoches*^42,43,45,46,50^, hence keeping internal relationships (and corresponding conflicts, see bellow) of the lineages unresolved. Serious phylogenetic incongruence between ‘nuc’ and ‘cp’ tree topologies of Catasetinae has been firstly identified by^63^, but a discussion about the respective plausibility of the trees has been remained undone.

All major clades in our ‘nuc’ phylogeny are consistent with morphological concepts of *Cycnoches* (Fig. 1a). *Cycnoches hagii* (Clade A) differs from other *Cycnoches* species by the fleshy oblong, curved pair set of calli located towards the base of the labellum. Clade B includes species with conspicuously large male flowers and an entire to 4-lobed labellum blade. Clade C comprises species with proportionately small male flowers and labella with 8–10 marginal projections^48^. The consistence between ‘nuc’ molecules and morphology is challenged by an apparent correlation between the ‘cp’ molecular tree and the distributions of particular lineages (Fig. 1b): Clade I comprises species occurring mostly in Amazonia, Caribbean-Llanos, and Guyana shield regions (except *C. warzsewiczii* present in Panama and Costa Rica: Fig. S4). Clade II, in turn, includes species exclusively distributed in Central America and Choco regions.

Topological incongruence between phylogenies derived from different DNA data partitions is a widespread phenomenon in phylogenetic inferences^64,65^. Examples from angiosperms (e.g., Araceae^37^, Asteraceae^66^, Saxifragaceae^67^) have long revealed similar conflicting patterns, in which the ‘nuc’ phylogeny is in accordance with morphology, whereas the ‘cp’ relationships correlate to geographical distributions. For the particular case of *Cycnoches*, hybridization might have explanatory power for the nuclear-plastid conflict observed in the clade. Euglossine-bee-pollinated orchids such as *Cycnoches* produce a blend of volatile compounds, which attracts male Euglossine bees, and pollination takes place while bees collect such compounds produced by specialized tissues in the flower^68^. Species-specific production of floral blends and therefore attraction of a unique set of pollinator(s) has been accounted as an isolative reproductive barrier in Euglossine bee pollinated orchids^41,69^. Nevertheless, intra-specific variation of fragrances produced by the flower has been reported in several orchid lineages such as *Stanhopea*^70^ and even *Cycnoches*^71^. Fragrance variation may result on attraction of a set of pollinators that are shared by species co-occurring in the same biogeographical region (e.g. *C. dianae, C. guttulatum*, and *C. pachydactylon*^71^; see also Fig. S4) with similar composition of the fragrance profile, providing an opportunity to hybridization to occur. Little is known about pollinators of *Cycnoches*, but our own observations of pollinator sharing between species and presence of polymorphic species may indicate past hybridization processes.

## Conclusion

Based on a solid, comprehensively sampled phylogeny we provide macroevolutionary evidence for a South American origin of *Cycnoches.* Our biogeographical analysis indicates colonization of Central America via a direct migration from the Amazonian basin. More importantly, the analyses support three recent trans-Andean, bidirectional migration processes between Central America and Amazonia, which is indicative for a minor effect of the Andean barrier on swan orchids migration. Consequently, our study enlightens the limited role of Andean mountain building on the range evolution and diversification of lowland Neotropical epiphytic lineages.

## Material and methods

### Taxon sampling, DNA sequencing and phylogenetic analyses

Species names, geographical origins, voucher specimens, and GenBank accession numbers of sequences included in phylogenetic analyses are provided in Table S1. Our study builds-up upon the DNA data matrices generated by^40,46,63^. Genomic DNA was extracted from herbarium and fresh leaf material with the NucleoSpin^®^ plant kit (Macherey-Nagel; Düren, Germany). We amplified and sequenced nuclear (consistently referred as ‘nuc’ henceforth) ribosomal external and internal transcribed spacers (ETS and ITS, respectively), and a fragment of the *Xdh* gene. We also sequenced a ~1500 bp fragment of the plastid (referred to as ‘cp’) gene *ycf*1, as well as the *trn*S–*trn*G intergenic spacer. Amplification and sequencing settings, as well as sequencing primers used for ITS, ETS, *Xdh*, *trn*S–*trn*G, and *ycf*1 are the same as in^63,72^ (Table S2). In this study, 84 sequences were newly generated (Table S1).

Loci were aligned separately using MAFFT 7.1^73^. For ‘nuc’ ribosomal RNA loci and ‘cp’ *trn*S–*trn*G spacer, secondary structure of molecules was taken into account (i.e., the-qINSi option). Congruence between ‘nuc’ and ‘cp’ datasets was assessed following Pérez-Escobar et al.^63^ and using PACo^74^. The procedure is available as a pipeline (http://www.uv.es/cophylpaco/) and was also employed to identify operational terminal units (OTUs) from the ‘cp’ dataset that conflicting with the ‘nuc’ dataset (potential outliers detected by PACo are shown in Figs S1–S2). A detailed explanation on PACo and a rationale on outlier handling is provided Extended Materials and Methods of Appendix S1.

Phylogenetic analyses of separate and concatenated loci were carried out under maximum likelihood (ML) and Bayesian inference (BI). The best-fitting evolutionary models for ML and Bayesian analyses (for each data partition) were selected using jModelTest v.2.1.6^75^, relying on a Likelihood Ratio Test (LRT) and the Akaike information criterion (AIC) (Table S3). Phylogenetic inference relied on the ML approach implemented in RAXML-HPC v.8.2.4^76^ and BI as implemented in MRBAYES v.3.2.2^77^ and were carried out on the CIPRES Science Gateway computing facility^78^. Bayesian inferences were performed with two independent runs, each with four Markov chains Monte Carlo (MCMC) running for 30 million generations each, and sampled every 1000^th^ generation (all other prior settings by default). Log files derived from MRBAYES were examined, and the convergence of MCMC was assessed using TRACER (available at: http://tree.bio.ed.ac.uk/software/tracer/). Node support values were assessed for both the ML tree using Maximum Likelihood Bootstrap Support (MLBS) and the consensus Bayesian tree using posterior probabilities (PP).

### Molecular clock dating

A few orchid macrofossils are available for Orchidaceae^79,80^, but these are assigned to lineages very distantly related to our groups of interest. Using distant outgroups to calibrate our *Cycnoches* phylogeny would have created extensive sampling heterogeneities, which can result in spurious results^81^. Thus, we had to rely on secondary calibrations. In order to obtain the best secondary calibration points possible, we first generated an Orchidaceae-wide fossil-calibrated phylogenies, sampling 316 orchid species sampled as evenly as possible along the tree. Loci, number of sequences and settings for absolute age estimation of the Orchidaceae-wide fossil calibrate phylogeny are provided in the Extened Materials and Methods of Appendix S1. The ages obtained were very similar to recent orchid dating studies^82,83^ and the dated phylogeny is shown in Figure S3.

We selected two secondary calibrations for the dating of *Cycnoches*: (*i*) the crown group of Catasetinae was set to 19.8 Mya with a standard deviation of 4 to reflect the 95% CI, and (*ii*) and the root of the *Cycnoches* tree (i.e., MRCA of *Cyrtopodium* + Catasetinae) was set to 27.1 Mya with a standard deviation of 6. To explore the clock-likeness of the data, we used both strict clock and uncorrelated lognormal clock models, and compared different tree priors (pure-birth, standard birth-death, and incomplete sampling birth-death). For strict molecular clock calibration, we placed a single constraint only at the tree root (27.1 Mya with a standard deviation of 6) using a normal distribution. The best-fitting tree speciation model was selected using Bayes factors calculated from marginal likelihoods computed for every model using the stepping-stone sampling^84^ (Table 1). For each clock model, we ran two MCMC analyses with 20 million generations each, sampled every 1000^th^ generation. For the relaxed molecular clock analyses, we estimated the coefficient of variation (CV) to inform us on the rate heterogeneity among branches (CV approaching 0 indicates that a strict clock model cannot be rejected). Parameter convergence was confirmed using TRACER (http://tree.bio.ed.ac.uk/software/tracer/). All dating analyses were performed at the CIPRES Science Gateway computing facility^78^.

### Ancestral area estimations

Species ranges of *Cycnoches* of were coded from the collection site and type locality of the material sequenced, which reflect the distribution ranges for every taxa included in our phylogeny (except by *C. chlorochilon* and *C. pentadactylon,* which also occur in Central America and Southeastern South America, respectively; see bellow). Distribution data of *Cycnoches* and outgroup taxa was obtained from own field observations, literature^85,86^ and from herbarium specimens (Herbaria AMES, COL, F, M, MO, SEL, US); this information was employed to code outgroup distribution ranges. Biogeographical areas were derived from distribution maps of the orchids under investigation (Fig. S4) as well as species distributions observed in other plant lineages (e.g., Rubiaceae^6^; Bromeliaceae^38^). We coded for eight biogeographical areas using the R-package ‘SpeciesGeocodeR’^87^: (1) Central America comprising southern Mexico through Panama; (2) Caribbean-Llanos comprising the coastal northernmost areas and plains of Colombia and Venezuela^88^; (3) Guiana Shield encompasses areas above 800 m in Colombia, Venezuela, Brazil, Guyana, Suriname, and French Guiana; (4) Amazonia encompassing lowlands and pre-montane forest below 800 m in Colombia, Ecuador, Peru, Brazil, Venezuela, Guyana, Suriname, and French Guiana^6^; (5) Chocó comprising lowlands below 500 m of the western Andes in Colombia and Ecuador; (6) Northern Andes including elevated areas above 800 m from Southernmost Peru to Northern Colombia and Northeast Venezuela; (7) Central Andes comprising areas above 800 m in Northern Peru to Northern Chile and Northeast Argentina; (8) South-eastern South America encompassing part of the Brazilian shield, the Atlantic forest, South-eastern Bolivia, Paraguay, Uruguay, and Northern Argentina. A map with the eight biogeographical areas, number of species for each area, and the corresponding distribution records of *Cycnoches* plus outgroups are provided in Figure S4.

To infer ancestral areas in *Cycnoches*, we used the R-package ‘BioGeoBEARS’ (Biogeography with Bayesian and Likelihood Evolutionary Analysis in R script^89^). Using BioGeoBEARS, we tested the fit of six biogeographic models with and without founder-event speciation (or jump speciation), altogether testing the role and contribution of evolutionary processes that were taken into account to explain today’s observed distributions (i.e., range expansions, local extinctions, founder-event speciation, vicariance, and speciation despite sympatry) in a joint statistical framework. It is therefore capable of model testing and of determining, which process fits better the geographical and phylogenetic data for any particular clade.

## Acknowledgements

We are grateful to Martina Silber (Munich, Germany) for valuable assistance in the lab, to Mario Blanco (San José, Costa Rica), Diego Bogarín (San José, Costa Rica), Norman Cash and family (Managua, Nicaragua) for the great help provided during field trip, to Boris Schlumpberger (Hannover, Germany), Andres Maduro (Panama City, Panama), Ivan G. Alvarez (David, Panama), Lankester Botanical Garden (Cartago, Costa Rica), Gustavo Romero (Herbarium HUH, Cambridge, U. S. A) and University of Costa Rica for providing plant material. We also thank the Smithsonian Tropical Research Institute (Panama City) and Costa Rican Ministry of Environment (MINAET) for providing research permits (No. SE/AP-20-13 and 026-2013-SINAC, respectively). We finally thank Alexandre Antonelli (Gothenburg, Sweden), for granting access to graphical material employed in Fig. 3. This work was supported by the Colombian Administrative Department of Science and Technology (COLCIENCIAS; granted to OAPE). We thank the Deutsche Forschungsgemeinschaft (granted to GG, GE 828/12-1) for financial support.

### Author contributions

O.A.P.E., M.G. and F.L.C. designed research; G.G. and O.A.P.E. collected samples; O.A.P.E. and E.P. performed all the lab work; O.A.P.E., F.L.C., and G.C. performed all analyses; O.A.P.E., M.G., F.C., G.C., G.G., B.K. and E.P. wrote the manuscript under the lead of O.A.P.E., M.G, and F.C.

### Competing financial interest

The authors declare no competing financial interest.

### Supplementary material

**Appendix S1.** Extended Materials and Methods, supplementary tables and figures.

## References

1. Humboldt, A. Voyage aux regions equinoxiales du Nouveau Continent. (N. Mazé, 1820).

2. Darwin, C. Geological observations of South America. (Smith, Elder and CO., 1846).

3. Jaramillo, C., Rueda, M. J. & Mora, G. Cenozoic plant diversity in the Neotropics. Science (80-.). 311, 1893–1896 (2006).

4. Antonelli, A. & Sanmartín, I. Why are there so many plant species in the Neotropics? Taxon 60, 403–414 (2011).

5. Myers, N., Mittermeier, R. A., Mittermeier, C. G., Fonseca, G. A. B. & Kent, J. Biodiversity hotspots for conservation priorities. Nature 403, 853–858 (2000).

6. Antonelli, A., Nylander, J. A. A., Persson, C. & Sanmartín, I. Tracing the impact of the Andean uplift on Neotropical plant evolution. Proc. Natl. Acad. Sci. U. S. A. 106, 9749–9754 (2009).

7. Hoorn, C. et al Amazonia Through Time: Andean uplift, climate change, landscape evolution, and biodiversity. Science (80-.). 330, 927–931 (2010).

8. Bacon, C. D. et al Biological evidence supports an early and complex emergence of the Isthmus of Panama. Proc. Natl. Acad. Sci. 112, 6110–6115 (2015).

9. Lagomarsino, L., Condamine, F. L., Antonelli, A., Mulch, A. & Davis, C. C. The abiotic and biotic drivers of rapid diversification in Andean bellflowers (Campanulaceae). New Phytol. 210, 1430–1432 (2016).

10. Hughes, C. E., Pennington, R. T. & Antonelli, A. Neotropical plant evolution: assembling the big picture. Bot. J. Linn. Soc. 171, 1–18 (2013).

11. Moonlight, P. W. et al Continental-scale diversification patterns in a megadiverse genus: the biogeography of Neotropical Begonia. J. Biogeogr. 42, 1–13 (2015).

12. Pennington, R. T. et al Contrasting plant diversification histories within the Andean biodiversity hotspot. Proc. Natl. Acad. Sci. 107, 13783–13787 (2010).

13. Luebert, F., Hilger, H. H. & Weigend, M. Diversification in the Andes: age and origins of South American Heliotropium lineages (Heliotropiaceae, Boraginales). Mol. Phylogenet. Evol. 61, 90–102 (2011).

14. Hughes, C. & Eastwood, R. Island radiation on a continental scale: exceptional rates of plant diversification after uplift of the Andes. Proc. Natl. Acad. Sci. U. S. 103, 10334–10339 (2006).

15. Nevado, B., Atchison, G. W., Hughes, C. E. & Filatov, D. A. Widespread adaptive evolution during repeated evolutionary radiations in New World lupins. Nat. Commun. 7, 1–9 (2016).

16. Bacon, C. D., Mora, A., Wagner, W. L. & Jaramillo, C. A. Testing geological models of evolution of the Isthmus of Panama in a phylogenetic framework. Bot. Linn. Soc. 171, 287–300 (2013).

17. Luebert, F. & Weigend, M. Phylogenetic insights into Andean plant diversification. 2, 1–17 (2014).

18. van der Hammen, T. The Pleistocene changes of vegetation and climate in Tropical South America. J. Biogeogr. 1, 3–26 (1974).

19. Martínez, C., Madriñán, S., Zavada, M. & Jaramillo, C. A. Tracing the fossil pollen record of Hedyosmum (Chloranthaceae), an old lineage with recent Neotropical diversification. Grana 52, 161–180 (2013).

20. Ghosh, P., Garzione, C. N. & Eiler, J. M. Rapid uplift of the Altiplano revealed through 13C-18O bonds in Paleosol carbonates. Science (80-.). 311, 511–515 (2006).

21. Kar, N. et al Rapid regional surface uplift of the northern Altiplano plateau revealed by multiproxy paleoclimate reconstruction. Earth Planet. Sci. Lett. 447, 33–47 (2016).

22. Hoorn, C., Guerrero, J., Sarmiento, G. A. & Lorente, M. A. Andean tectonics as a cause for changing drainage patterns in Miocene northern South America. Geology 23, 237–240 (1995).

23. Gregory-Wodzicki, K. M. Uplift history of the Central and Northern Andes: a review. Geol. Soc. Am. Bull. 112, 1091–1105 (2000).

24. Armijo, R., Lacassin, R., Coudurier-Curveur, A. & Carrizo, D. Coupled tectonic evolution of Andean orogeny and global climate. Earth Sci. Rev. 143, 1–35 (2015).

25. Gentry, A. H. Neotropical floristic diversity: phytogeographical connections between Central and South America, Pleistocene climatic fluctuations, or an accident of the Andean orogeny? Ann. Missouri Bot. Gard. 69, 557–593 (1982).

26. Richter, M., Diertl, K., Emck, P., Peters, T. & Beck, E. Reasons for an outstanding plant diversity in the tropical Andes of Southern Ecuador. Landsc. Online 12, 1–35 (2009).

27. Pirie, M. D., Chatrou, L. W., Mols, J. B., Erkens, R. H. J. & Oosterhof, J. ‘Andean-centred’ genera in the short-branch clade of Annonaceae: Testing biogeographical hypotheses using phylogeny reconstruction and molecular dating. J. Biogeogr. 33, 31–46 (2006).

28. Janzen, D. H. & Martin, P. S. Neotropical anachronisms: the fruits the gomphotheres ate. Science (80-.). 215, 19–27 (1982).

29. Uribe-Convers, S. & Tank, D. C. Shifts in diversification rates linked to biogeographic movement into new areas: An example of a recent radiation in the Andes. Am. J. Bot. 102, 1–16 (2015).

30. Sánchez-Baracaldo, P. Phylogenetics and biogegraphy of the Neotropical fern genera Jamesonia and Eriosorus (Pteridaceae). Am. J. Bot. 91, 274–284 (2004).

31. Kreft, H., Koster, N., Kuper, W., Nieder, J. & Barthlott, W. Diversity and biogeography of vascular epiphytes in Western Amazonia, Yasuni, Ecuador. J. Biogeogr. 31, 1463–1476 (2004).

32. Krömer, T., Kessler, M., Gradstein, S. R. & Acebey, A. Diversity patterns of vascular epiphytes along an elevational gradient in the Andes. J. Biogeogr. 32, 1799–1809 (2005).

33. Gentry, A. H. & Dodson, C. H. Diversity and biogeography of Neotropical vascular epiphytes. Ann. Missouri Bot. Gard. 74, 205–233 (1987).

34. Mutke, J. & Barthlott, W. Patterns of vascular plant diversity at continental to global scales. Biol. Skr. 55, 521–531 (2005).

35. Kier, G. et al Global patterns of plant diversity and floristic knowledge. J. Biogeogr. 32, 1107–1116 (2005).

36. Zotz, G. The systematic distribution of vascular epiphytes-a critical update. Bot. J. Linn. Soc. 171, 453–481 (2013).

37. Nauheimer, L., Boyce, P. C. & Renner, S. S. Giant taro and its relatives: a phylogeny of the large genus Alocasia (Araceae) sheds light on Miocene floristic exchange in the Malesian region. Mol. Phylogenet. Evol. 63, 43–51 (2012).

38. Givnish, T. J. et al Phylogeny, adaptive radiation, and historical biogeography in Bromeliaceae: Insights from an eight-locus plastid phylogeny. Am. J. Bot. 98, 872–895 (2011).

39. Pridgeon, A. M., Cribb, P. J., Chase, M. W. & Rasmussen, F. N. Genera Orchidacearum: Vol. 5. Epidendroideae (part two). (Oxford University Press, 2009).

40. Pérez-Escobar, O. A., Gottschling, M., Whitten, W. M., Salazar, G. & Gerlach, G. Sex and the Catasetinae (Darwin’s favourite orchids). Mol. Phylogenet. Evol. 97, 1–10 (2016).

41. Ramirez, S. R. et al Asynchronous Diversification in a Specialized Plant-Pollinator Mutualism. Science (80-.). 333, 1742–1746 (2011).

42. Chase, M. W. & Pippen, J. S. Seed morphology and phylogeny in Subtribe Catasetinae (Orchidaceae). Lindleyana 5, 126–133 (1990).

43. Romero, G. A. Phylogenetic relationships in Subtribe Catasetinae (Orchidaceae, Cymbidieae). Lindleyana 5, 160–181 (1990).

44. Stern, W. & Judd, W. Comparative anatomy and systematics of Catasetinae (Orchidaceae). Bot. J. Linn. Soc. 136, 153–178 (2001).

45. Batista, J. N. et al Molecular phylogenetics of Neotropical Cyanaeorchis (Cymbidieae, Epidendroideae, Orchidaceae): Geographical rather than morphological similarities plus a new specie. Phytotaxa 156, 251–272 (2014).

46. Whitten, W. M., Neubig, K. M. & Williams, N. H. Generic and subtribal relationships in neotropical Cymbidieae (Orchidaceae) based on matK/ycf1 plastid data. Lankesteriana 13, 375–392 (2014).

47. Carr, G. F. J. The genus Cycnoches: species and hybrids. Orchid Rev. 1–31 (2012).

48. Gerlach, G. & Pérez-Escobar, O. A. Looking for missins swans: Phylogenetics of Cycnoches. Orchids 83, 434–437 (2014).

49. Darwin, C. On the various contrivances by which british and foreign orchids are fertilised by insects. (Appleton and CO., 1877).

50. Pridgeon, A. M. & Chase, M. W. in Proceeding of the 15th World Orchid Conference 275–281 (1998).

51. Romero, G. A. & Gerlach, G. in Flora Mesoamericana (Missouri Botanical Garden Press).

52. Garzione, C. N., Molnar, P., Libarkin, J. C. & Macfadden, B. J. Rapid late Miocene rise of the Bolivian Altiplano: evidence for removal of mantle lithosphere. Earth Planet. Sci. Lett. 241, 543–556 (2006).

53. Garzione, C. N. et al Rise of the Andes. Nature 320, 1304–1307 (2008).

54. Garzione, C. N. et al Clumped isotope evidence for diachronous surface cooling of the Altiplano and pulsed surface uplift of the Central Andes. Earth Planet. Sci. Lett. 393, 173–181 (2014).

55. Sánchez-Baracaldo, P. & Thomas, G. H. Adaptation and convergent evolution within the Jamesonia-Eriosorus complex in high-elevation biodiverse andean hotspots. PLoS One 9, e110618 (2014).

56. Arditti, J. & Ghani, A. K. A. Numerical and physical properties of orchid seeds and their biological implications. New Phytol. 145, 367–421 (2000).

57. Jersáková, J. & Malinová, T. Spatial aspects of seed dispersal and seedling recruitment in orchids. New Phytol. 176, 237–241 (2004).

58. Barthlott, W., Große-Veldmann, B. & Korotkova, N. Orchid seed diversity: a scanning electron microscopy survey. (Botanic Garden and Botanical Museum Berlin, 2014).

59. acquemyn, H. et al A spatially explicit analysis of seedling recruitment in the terrestrial orchid Orchis purpurea. New Phytol. 176, 448–459 (2004).

60. Close, R. C., Moar, N. T., Tomlinson, A. I. & Lowe, A. D. Aerial dispersal of biological material from Australia to New Zealand. Int. J. Biometeorol. 22, 1–19 (1978).

61. Gandawijaja, D. & Arditti, J. The orchids of Krakatau: evidence for a mode of transport. Ann. Bot. 52, 127–130 (1983).

62. Nakamura, S. J. & Hamada, M. On the seed dispersal of an achlorophyllous orchid, Galeola septentrionalis. J. Japanese Bot. 53, 260–263 (1978).

63. Pérez-Escobar, O. A., Balbuena, J. A. & Gottschling, M. Rumbling orchids: How to assess divergent evolution between chloroplast endosymbionts and the nuclear host. Syst. Biol. 65, 51–65 (2016).

64. Rokas, A., Williams, B. L., King, N. & Carroll, S. B. Genome-scale approaches to resolving incongruence in molecular phylogenies. Nature 425, 798–804 (2003).

65. Salichos, L., Stamatakis, A. & Rokas, A. Novel information theory-based measures for quantifying incongruence among phylogenetic trees. Mol. Biol. Evol. 31, 1261–1271 (2014).

66. Rieseberg, L. H., Beckstrom-Sternberg, S. & Doan, K. Helianthus annuus ssp. texanus has chloroplast DNA and nuclear ribosomal RNA genes of Helianthus debilis ssp. cucumerifolius. Proc. Natl. Acad. Sci. U. S. A. 87, 593–597 (1990).

67. Rieseberg, L. H. & Soltis, D. E. Phylogenetic consequences of cytoplasmatic gene flow in plants. Evol. Trends Plants 5, 65–84 (1991).

68. Gerlach, G. & Schill, R. Composition of Orchid Scents Attracting Euglossine Bees. Bot. Acta 104, 379–384 (1991).

69. Dressler, R. L. Pollination by Euglossine Bees. Evolution (N. Y). 22, 202–210 (1967).

70. Williams, H. & Whitten, W. M. Orchid floral fragrances and male euglossine bees: methods and advances in the last sesquidecade. Biol. Bull. 164, 355–395 (1983).

71. Gregg, K. B. Variation in floral fragrances and morphology: incipient speciation in Cycnoches? Bot. Gaz. 144, 566–576 (1983).

72. Irimia, R.-E., A. Pérez-Escobar, O. & Gottschling, M. Strong biogeographic signal in the phylogenetic relationships of Rochefortia Sw. (Ehretiaceae, Boraginales). Plant Syst. Evol. 301, 1509–1516 (2014).

73. Katoh, K. & Standley, D. M. MAFFT multiple sequence alignment software version 7: improvements in performance and usability. Mol. Biol. Evol. 30, 772–780 (2013).

74. Balbuena, J. A., Míguez-Lozano, R. & Blasco-Costa, I. PACo: a novel procrustes application to cophylogenetic analysis. PLoS One 8, e61048 (2013).

75. Darriba, D., Taboada, G. L., Doallo, R. & Posada, D. jModelTest 2: more models, new heuristics and parallel computing. Nat. Methods 9, 772–772 (2012).

76. Stamatakis, A. RAxML version 8: A tool for phylogenetic analysis and post-analysis of large phylogenies. Bioinformatics 30, 1312–1313 (2014).

77. Ronquist, F. et al Mrbayes 3.2: Efficient Bayesian phylogenetic inference and model choice across a large model space. Syst. Biol. 61, 539–542 (2012).

78. Miller, M. A. et al A RESTful API for access to phylogenetic tools via the CIPRES Science Gateway. Evol. Bioinforma. 11, 43–48 (2015).

79. Ramírez, S. R., Gravendeel, B., Singer, R. B., Marshall, C. R. & Pierce, N. E. Dating the origin of the Orchidaceae from a fossil orchid with its pollinator. Nature 448, 1042–1045 (2007).

80. Conran, J. G., Bannister, J. M. & Lee, D. E. Earliest orchid macrofossils: Early Miocene Dendrobium and Earina (Orchidaceae: Epidendroideae) from New Zealand. Am. J. Bot. 96, 466–474 (2009).

81. Drummond, A. J. & Bouckaert, R. R. Bayesian evolutionary analysis with BEAST 2. (Cambridge University Press, 2014).

82. Chomicki, G. et al The velamen protects photosynthetic orchid roots against UV-B damage, and a large dated phylogeny implies multiple gains and losses of this function during the Cenozoic. New Phytol. 205, 1330–1341 (2014).

83. Givnish, T. J. et al Orchid historical biogeography, diversification, Antarctica and the paradox of orchid dispersal. 1–12 (2016). doi:10.1111/jbi.12854

84. Xie, W., Lewis, P. O., Fan, Y., Kuo, L. & Chen, M. H. Improving marginal likelihood estimation for bayesian phylogenetic model selection. Syst. Biol. 60, 150–160 (2011).

85. Carr, G. F. J. The genus Cycnoches and its species Part 4: modern studies of pollinators. Orchid Rev. 221–225 (2006).

86. Romero, G. A. in Genera Orchidacearum: Vol. 5. Epidendroideae (part two) (eds. Pridgeon, A. M., Cribb, P. J., Chase, M. W. & Rasmussen, F. N.) 11–40 (Oxford University Press, 2009).

87. Töpel, M. et al SpeciesGeoCoder: Fast categorisation of species occurrences for analyses of biodiversity, biogeography, ecology and evolution. BioRxiv 9274, 1–15 (2014).

88. Morrone, J. J. Biogeographic areas and transition zones of Latin America and the Caribbean islands based on panbiogeographic and cladistic analyses of the entomofauna. Annu. Rev. Entomol. 51, 467–494 (2006).

89. Matzke, N. J. Model selection in historical biogeography reveals that founder-event speciation is a crucial process in island clades. Syst. Biol. 63, syu056–(2014).

